# Martial arts training is related to implicit intermanual transfer of visuomotor adaptation

**DOI:** 10.1101/2020.06.09.141887

**Authors:** Susen Werner, Koki Hasegawa, Kazuyuki Kanosue, Heiko K. Strüder, Tobias Göb, Tobias Vogt

## Abstract

Recent work identified an explicit and implicit transfer of motor learning with one limb to the other untrained limb. Here we pursue the idea that different individual factors differently contribute to the amount of explicit and implicit intermanual transfer of sensorimotor adaptation. In particular we tested a group of judo athletes who show enhanced right-hemispheric involvement in motor control and a group of equally trained athletes as control participants. After adaptation to a 60° visual rotation, we estimated awareness of the perturbation and transfer to the untrained, non-dominant left hand in two experiments. We measured the total amount of intermanual transfer (explicit plus implicit) by telling the participants to repeat what was learned during adaptation and the amount of implicit transfer by instructing the participants to refrain from using what was learned but to perform movements as during baseline. We found no difference between the total intermanual transfer of judokas and running experts, with mean absolute transfer values of 42.4° and 47.0°. Implicit intermanual transfer was very limited but larger in judo than in general sports athletes, with mean values of 5.2° and 1.6°. A multiple linear regression analysis further revealed that total intermanual transfer, which largely represents the explicit transfer, is related to awareness of the perturbation while implicit intermanual transfer can be predicted by judo training, amount of total training, speed of adaptation and handedness scores. The findings are discussed in relation to neuronal mechanisms such as hemispheric interactions and functional specialization underlying intermanual transfer of motor learning.

## 1. Introduction

There are good reasons for studying the generalization of motor learning from one arm to the other, untrained arm. On a behavioral level, understanding the mechanisms of intermanual transfer can help athletes and coaches design effective training measures (Haaland & Hoff, 2003) or it can be useful in upper-limb prosthesis training (Romkema *et al.*, 2017). On a neuronal level, the study of intermanual transfer can broaden our knowledge about hemispheric interactions (Chase & Seidler, 2008) or it can unveil the global and local nature of internal representations of motor learning (Harris, 1963; Lefumat *et al.*, 2015).

So far several internal or individual factors determining the amount of intermanual transfer have been identified. First, Chase and Seidler (2008) reported an increased intermanual transfer of sensorimotor adaptation in less strongly left-handed and better intermanual transfer of sequence learning in less strongly right-handed participants. Second, participants who adapt faster and, thus, spend more time at plateau, should also display an increased amount of transfer, since increased time at adaptation plateau also positively effects intermanual transfer (Block & Celnik, 2013). This notion is supported by the fact that a longer adaptation training schedule resulted in more time spend at plateau and substantially increased retention of intermanual transfer (Joiner *et al.*, 2013). A third factor determining intermanual transfer is related to the explicit and implicit process of sensorimotor adaptation. Recent works have demonstrated a relation between awareness of the specific pattern of perturbation and intermanual transfer (Malfait & Ostry, 2004; Poh *et al.*, 2016; Werner *et al.*, 2019). In addition, variability during learning phase, which is also related to the explicit process of adaptation, was further identified to enhance intermanual transfer of adaptation to a Coriolis force (Lefumat *et al.*, 2015). In fact, Poh et al. (2016) recently suggested that explicit and implicit processes are not restricted to adaptation itself but that intermanual transfer also comprises of explicit and implicit components. These authors measured the amount of explicit intermanual transfer by assessing verbal reports of subjects’ aiming direction before each reaching movement. Their results suggest complete transfer of the explicit but only incomplete transfer of the implicit process. However, it remains unclear if the above mentioned individual factors (handedness, speed of adaptation, explicit/implicit processes) contribute to the explicit or the implicit intermanual transfer of sensorimotor adaptation. In the present project we aimed at scrutinizing their involvement in young healthy athletes as well as in martial artists, a group of highly specialized participants.

Martial arts training is characterized by complex coordinate movement of both sides (Dopico, Iglesias-Soler, *et al.*, 2014). In striking sports both arms and legs are involved (Layton, 1993; Neto *et al.*, 2012; Maeda *et al.*, 2014) and also in grabbling sports most techniques involve bimanual actions (Mikheev *et al.*, 2002). Nevertheless most martial artists have their preferred stand position with one foot in front of the other (Mikheev *et al.*, 2002; Baker & Schorer, 2013) and their preferred side when performing a technique (Mayo *et al.*, 2019). However, martial arts athletes who can skillfully use both sides are more versatile and can choose between more options for action than fighters with strong side dominance (Dopico, Iglesias-soler, *et al.*, 2014; Mayo *et al.*, 2019). It is, thus, not surprising that highly experienced *kung fu* athletes show a lower strength of hand preference measured by the Edinburgh handedness inventory (Oldfield, 1971) than *kung fu* novices (Maeda *et al.*, 2014). However, a different picture emerges in *judo* athletes: First, Dopico et al. (Dopico, Iglesias-soler, *et al.*, 2014) found no relation between the preferred limb use in a variety of daily life tasks (motor dominance) and preferred side during execution of judo techniques (functional dominance) in judokas. According to that, general lateral preference of everyday activities is not associated with lateral preference in judo. This is in line with the notion, that handedness scores such as those measured by the handedness inventory only assess a fraction of whole body side preference and even hand preference is a multi-dimensional trait (Healey *et al.*, 1986; Steenhuis & Bryden, 1989). Second, Mikheev et al. (Mikheev *et al.*, 2002) found a dissociation of handedness scores and hemispheric lateralization in judo athletes. The authors investigated lateralization profiles in judokas by a variety of tasks, including lateralized visual pattern and verbal listening tasks, and found enhanced right-hemispheric involvement in motor control relative to control participants. All tested judokas were overall right-handed according to handedness scores. Therefore, the increased use of the non-dominant left hand in right-handed judo experts goes hand in hand with a decrease of hemispheric lateralization. In the present study we, thus, included a group of right-handed judo athletes in order to find out if their increased right-hemispheric involvement contributes to explicit or implicit intermanual transfer of visuomotor adaptation.

Our current study pursued the idea that different individual factors differently contribute to the amount of explicit and implicit intermanual transfer. All participants initially performed a visuomotor adaptation task with their dominant right arm. We then tested how well this task was transferred to the non-dominant, untrained, left arm. We measured individual factors such as handedness scores, speed of adaptation, awareness of the perturbation and amount of training and set them in relation to intermanual transfer. In a first experiment we investigated the total amount of intermanual transfer (explicit plus implicit) and in a second experiment we analyzed the amount of implicit intermanual transfer.

## 2. Materials and Methods

### 2.1 Participants

A total of 53 subjects participated in the study, all performing the task under the same experimental protocol. Their ages ranged from 18 to 38 years and they were all adults in the country of testing. Participants were age- and gender-matched between the experimental groups. The participants regular athletic activity was assessed by a questionnaire and handedness scores were measured by the Edinburgh inventory (Oldfield, 1971). None of the subjects had any experience in visuomotor adaptation research or exhibited overt sensorimotor deficits besides corrected vision. The experimental protocol was conducted according to the principles expressed in the Declaration of Helsinki and was pre-approved by the ethical committee of the German Sport University Cologne. All participants gave written informed consent.

#### 2.1.1 Experiment on total intermanual transfer

This experiment was designed to examine the effect of intensive judo and running training on visuomotor adaptation and total intermanual transfer. Two groups participated: The group of judo experts all had a minimum of five years of training (n = 12, age 20 ± 2, all male). The group of running experts also had a minimum of five years of training of a running discipline and, in addition, a maximum of six months training of any martial arts (n = 12, age 19 ± 1, all male). All participants competed at least on a national level and were part of the college team of the Waseda University, Tokyo, Japan. This first experiment was conducted at the Faculty of Sport Sciences of the Waseda University.

#### 2.1.2 Experiment on implicit intermanual transfer

This experiment was designed to examine the effect of judo and more general sports training on visuomotor adaptation and implicit intermanual transfer. Two groups participated in this experiment: The group of judo athletes had a minimum of four years of training (n = 13, age 26 ± 6, 3 female / 10 male). The group of sports experts also had a minimum of four years of training of a sports discipline (n = 16, age 26 ± 5, 4 female/12 male). Their sports included ball sports (soccer, volleyball, handball), racket sports (tennis, badminton), individual sports (apparatus gymnastics, track and field, swimming, triathlon, running, climbing), carnival sports (majorette) as well as general strength, coordination and fitness training. In order to compare judo training to general sports training we did not exclude any specific sports in this second group except for martial arts training. This second experiment was conducted at the Institute of Movement and Neurosciences of the German Sport University, Cologne, Germany.

### 2.2 Task

Subjects sat in front of the set up and used a pen to move on a digitizing tablet (Werner *et al.*, 2019). These movements were recorded and displayed as a light-blue cursor on a computer screen. The subjects received visual feedback of their movements by watching the screen through a mirror. The projected computer-generated image, thus, appeared in the plane in which the participants performed their movements, while the mirror prevented vision of the arm. A central dot and one of eight possible target dots were alternately displayed on the screen in addition to the cursor. The targets were presented in random order and all dots were yellow and 5 mm in diameter. The target dots were evenly distributed on an imaginary circle around the center with a radius of 5 cm and were displayed for a duration of 1000 ms. The central dot was displayed for a duration of 3 s plus a random interval of up to 500 ms. Participants were instructed to move the pen as precisely and as quickly as possible from the center to the lit target dot and back. Subjects performed their reaching movements during episodes of 35 s duration which were interrupted by rest breaks of 5 s. Approximately eight trails were performed during each episode.

### 2.3 Experimental design

Table 1 gives an overview of the experimental protocol. After *familiarization* with veridical feedback all participants conducted *baseline* episodes without visual feedback, i.e. no cursor visible, as well as baseline episodes with the left and right hand. In the following *adaptation* phase of 25 episodes, the visuomotor adaptation was induced by rotation of the cursor of 60 ° CCW around the central dot. Next came two episodes each of *inclusion* and *exclusion* to test for awareness in a process dissociation procedure as in Jacoby (1991) and Werner et al. (2015). This method is based on defining conscious knowledge as controllable knowledge. Thus, aware and unaware learning can be estimated by comparing performance when participants attempt to either express or repress a learned behavior. Prior to inclusion, subjects were, thus, instructed to ‘use what was learned during adaptation’ and prior to exclusion subjects were asked to ‘refrain from using what was learned, perform movements as during baseline’ (Werner *et al.*, 2015, 2019; Neville & Cressman, 2018). The order of inclusion and exclusion episodes was randomized between participants and no visual feedback was given in those episodes. During adaptation, inclusion, exclusion and intermitted *refresh* phases under rotated feedback participants used their right hand. In the subsequent test of *intermanual transfer*, movements were performed with the left hand without visual feedback. Visual feedback was not given in order to prevent confounding transfer with learning benefits to opposite limb learning (Joiner *et al.*, 2013; Poh *et al.*, 2016).

**Tab. 1:**
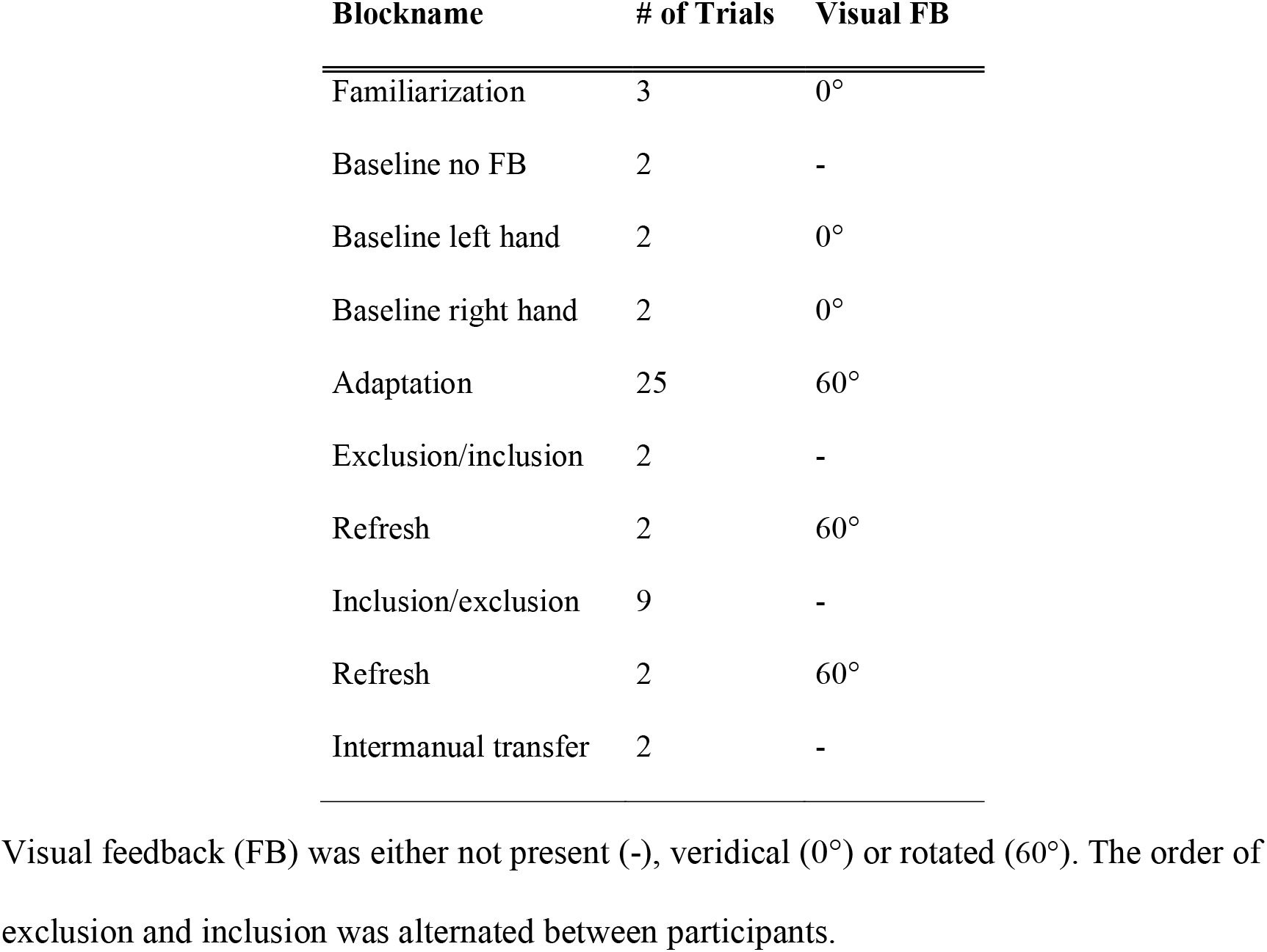
Experimental protocol.

| **Blockname** | **# of Trials** | **Visual FB** |
| --- | --- | --- |
| Familiarization | 3 | 0° |
| Baseline no FB | 2 | - |
| Baseline left hand | 2 | 0° |
| Baseline right hand | 2 | 0° |
| Adaptation | 25 | 60° |
| Exclusion/inclusion | 2 | - |
| Refresh | 2 | 60° |
| Inclusion/exclusion | 9 | - |
| Refresh | 2 | 60° |
| Intermanual transfer | 2 | - |

Visual feedback (FB) was either not present (-), veridical (0°) or rotated (60°). The order of exclusion and inclusion was alternated between participants.

The main difference between our two experiments was that we measured different components of intermanual transfer. In the first experiment we tested for the total amount of transfer, i.e. the explicit plus the implicit component. Therefore, participants were instructed to ‘repeat what was learned during adaptation’. In the second experiment we solely tested for the implicit amount of intermanual transfer. Here, participants were instructed to ‘refrain from using what was learned, perform movements as during baseline’. This way of probing for explicit and implicit components is in accordance with the inclusion and exclusion condition of the process dissociation procedure, respectively.

### 2.4 Data processing

From the questionnaires data we calculated the amount of total training as

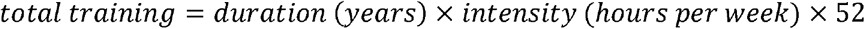

and a laterality quotient as

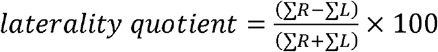

where ΣR (ΣL) is the sum of all right (left) items (Oldfield, 1971; Mikheev *et al.*, 2002). A quotient greater or less than +15% or −15% is interpreted as right-handedness or left-handedness; values between −15 and +15% mean both-handedness. A factor analysis of various items revealed that, in particular, those activities that occur in the personal space (such as combing hair as opposed to throwing a ball) are performed with the left hand in right-handed martial arts experts (Mikheev *et al.*, 2002). Therefore, we calculate another laterality quotient using only those items that take place in personal space. This corresponds to ‘Toothbrush’ and ‘Spoon’ in the first experiment (10-item Japanese version of the Edinburgh inventory) and to ‘Comb’, ‘Toothbrush’ and ‘Spoon’ in the second experiment (20-item English version of the Edinburgh inventory).

We further quantified participants’ reaching performance for each trial as *movement direction* with respect to the target direction 150 ms after movement onset, i.e., before feedback-based corrections could become effective (e.g. Prablanc and Martin 1992; Cressman et al. 2006). Movement onset was defined as a speed of 30 mm/s. From single trial data we calculated mean movement directions for each participant and episode. Group averages were used to visualize performance during baseline and adaptation. To determine the *speed of adaptation* we calculated the number of episode in which the participant first reached a plateau. This plateau was defined as mean of the participant’s last five episodes minus two times the standard deviation of all last five episodes of all participants. Therefore, speed of adaptation is greater if the number of plateau is smaller.

*Intermanual transfer* values were further calculated as mean movement directions of both transfer episodes. As done previously, we defined the amount of intermanual transfer as absolute values of transfer episodes (Sainburg and Wang 2002). Significant intermanual transfer shows as negative movement directions in the data. For better understanding we multiplied this parameter by −1, so that an increase in movement direction corresponds to an increase in intermanual transfer.

Moreover, we determined the amount of *awareness* from movement directions during exclusion and inclusion episodes. We first calculated exclusion and inclusion indices as

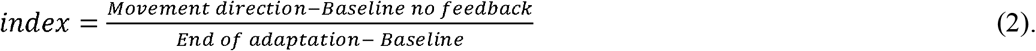

*Movement direction* is the mean movement direction of both exclusion or inclusion episodes for exclusion and inclusion indices, respectively. *Baseline* or *Baseline no feedback* were taken from all baseline episodes of the right hand with or without visual feedback, respectively, while *End of adaptation* was the movement direction of the last episode of adaptation phase. Following the logic of the process dissociation procedure, awareness can be estimated as the difference between exclusion and inclusion performance. Consequently, we determined the awareness index as inclusion minus exclusion index (Werner *et al.*, 2015, 2019).

For statistical analysis of group comparisons we performed t-tests in case of normality or Mann-Whitney U-tests if data did not follow a normal distribution or included significant outliners. Effect size is reported for significant differences as Cohen’s *d* or as Pearson correlation coefficient *r*, respectively. Normality within each group was explored by Kolmogorov-Smirnov test. Baseline and adaptation data were submitted to two analyses of variance (ANOVA) with the between-factor group (judo, running/sport) and the within-factor episode. Greenhouse-Geisser-adjustments were applied when necessary to compensate for heterogeneity of variances. Effect size is reported for significant differences as Eta-squared *η*^*2*^. In addition, multiple linear regressions were calculated with the dependent variable intermanual transfer and the independent variables group, speed of learning, awareness, total training and laterality quotient. Auto-correlation was explored by Durbin-Watson statistic. We checked for outliers using casewise diagnostics and studentized deleted residuals. In addition we tested for leverage points and checked their influence using Cook’s distance. In case of significant outliners or influential points we omitted this data point and repeated the analysis. All these statistical comparisons were performed using SPSS (Version 26.0. Armonk, NY: IBM Corp.).

## 3 Results

### 3.1 Adaptation

Figure 1 shows angular movement directions of all experimental phases and all participants of the first (Fig. 1A) and the second experiment (Fig. 1B). Data is provided as supporting information S1 on OSF (https://osf.io/khgzs/). Note that exclusion is depicted here before inclusion although, in fact, their order was randomized between participants. We can observe similar reaching directions for all participants throughout all experimental phases. For experiment one, movement directions of the running experts seem more variable than those of the judo experts in the beginning of adaptation. Movement directions of the final intermanual transfer phase come close to final adaptation level, indicating a large amount of total transfer. For experiment two, one participant from the judo group did not show any adaptation at all. Therefore, this data was excluded from all further analyses. Movement directions during test of intermanual transfer show very little intermanual transfer compared to experiment one.

**Fig. 1:**
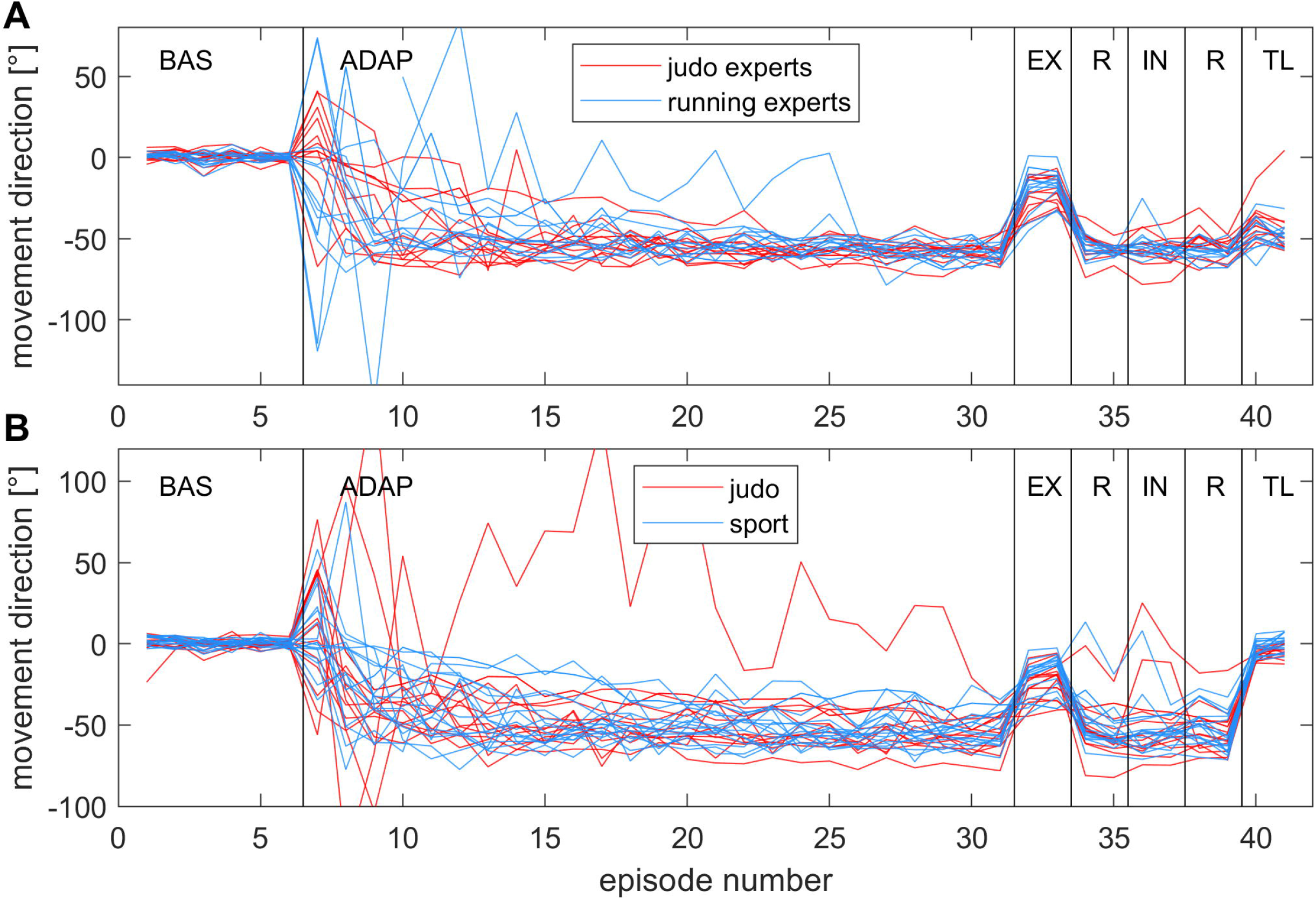
Individual movement directions of all experimental phases. Shown are angular movement directions with respect to target direction of all experimental phases and all participants of the first (A) and the second experiment (B). Lines indicate within-episode means for each judoka, running expert or sport athlete, respectively. Baseline (BAS), adaptation (ADAP), exclusion (EX), refresh (R), inclusion (IN) and intermanual transfer (TL) phase are depicted. Note that exclusion and inclusion order was randomized between participants.

Figure 2 illustrates mean movement directions for baseline and adaptation phases for both groups and experiments. All mean movement directions are close to zero during baseline in the first (Fig. 2A) as well as in the second experiment (Fig. 2B). During adaptation phase all groups show clear learning curves and movement directions close to −60° at the end of adaptation. Figure 2A shows that mean movement directions of the judo experts seem to reach a plateau earlier than those of the running experts in the first experiment whereas Figure 2B reveals no differences between learning curves of judo or general sports athletes in the second experiment. According to these observations, ANOVAs of baseline data revealed no significant effects for experiment one [group: F(1,22) = 0.081, p = 0.779; episode: F(3,60) = 2.76, p = 0.055; group × episode: F(3,60) = 0.51, p = 0.663] and two [group: F(1,26) = 2.23, p = 0.148; episode: F(3,85) = 0.65, p = 0.600; group × episode: F(3,85) = 1.51, p = 0.216]. Moreover, statistical analysis of adaptation data of the first experiment yielded an effect of episode [F(24,480) = 11.96, p < 0.001, *η*^*2*^ = 0.375] but not of group [F(1,20) = 0.30, p = 0.590]. The interaction between group and episode was marginally significant here [F(24,480) = 1.51, p = 0.058]. In addition, ANOVA of adaptation data of the second experiment revealed an effect of episode [F(4,110) = 23.84, p < 0.001, *η*^*2*^ = 0.478] but not of group [F(1,26) = 0.46, p=0.503] or of group × episode [F(4,110) = 0.60, p = 0.675]. Thus, all four experimental groups adapted to the visual perturbation, the running experts of experiment one seemed to adapt slightly slower than the judo experts, but final adaptation level did not differ between groups within experiments.

**Fig. 2:**
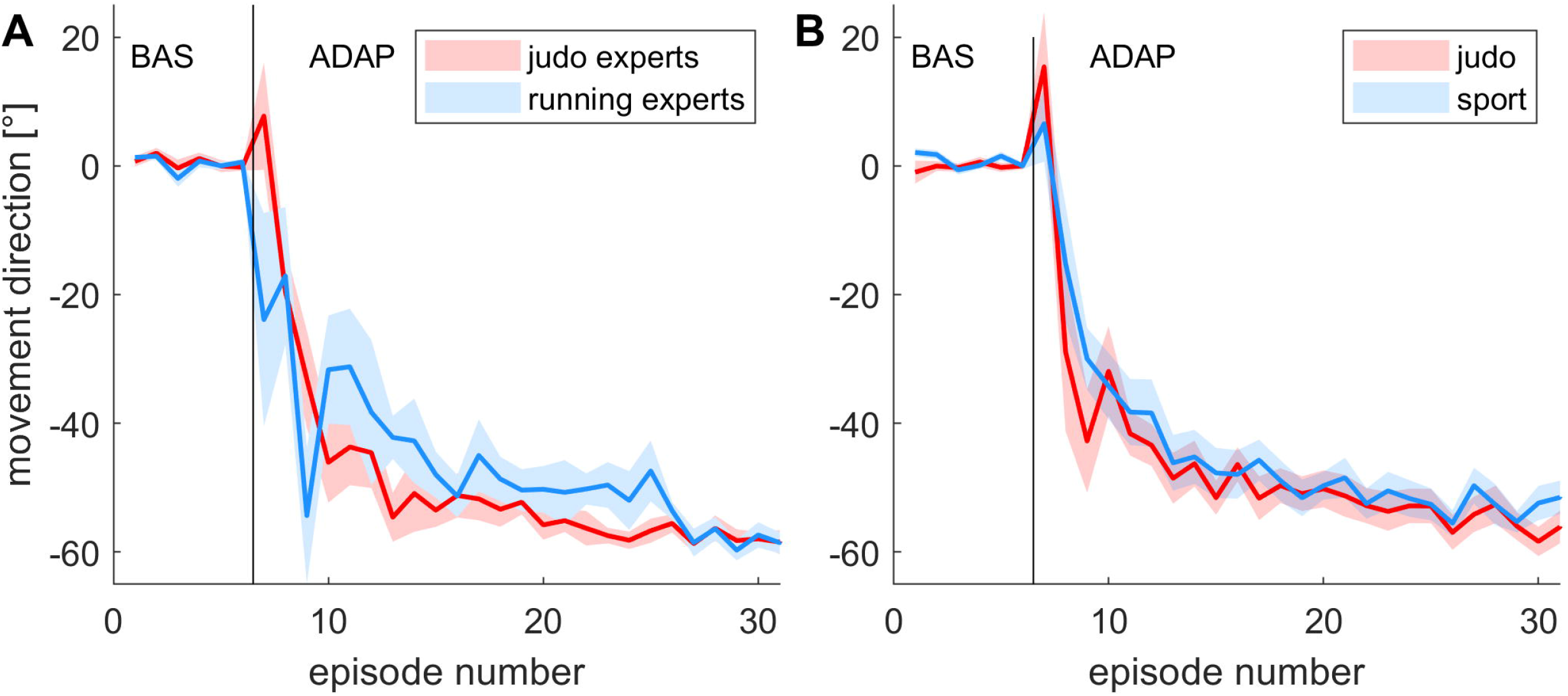
Mean movement directions of both groups. Shown are movement directions of baseline (BAS) and adaptation (ADAP). Lines indicate across-subject means, and the shaded area display standard errors. Judo and running group of the first (A) and judo and sport group of the second experiment (B) are depicted.

### 3.2 Total intermanual transfer and individual factors

Mean values of total intermanual transfer and individual factors of judo and running experts of the first experiment are depicted in Figure 3. One participant did not fill out one question in the activity questionnaire for assessment of the amount of total training and had to be excluded from the corresponding analyzes. Questionnaire data is provided as supporting information S2 on OSF (https://osf.io/khgzs/). Both groups show similar values for the total amount of intermanual transfer. Accordingly, group comparison revealed no significant difference [judo experts 42.4 ± 13.3°; running experts 47.0 ± 6.9°; U = −75, p = 0.478]. Figure 3 also depicts similar group levels for the individual factors awareness, total training and laterality quotient. Speed of adaptation is determined by the number of episode reaching plateau. Thus, a smaller number indicates larger speed of adaptation and, from Figure 3, judo experts seem to adapt faster than running experts. But still statistical analyzes of individual factors yielded neither a significant group effect for speed of adaptation [judo experts 8.6 ± 4.9, running experts 12.3 ± 6.6; t(22) = −1.580, p = 0.128] nor for awareness [judo experts 0.62 ± 0.23; running experts 0.64 ± 0.27; U = −0.12, p = 0.932], total training [judo experts 10929 ± 2680h; running experts 9776 ± 3480h; t(21) = 0.89, p = 0.381] or for laterality quotient [judo experts 81 ± 14; running experts 82 ± 15; t(22) = 0.18, p = 0.857]. Yet, Edinburgh inventory data showed increased both-handedness for items within personal space such as combing or brushing teeth in the judo group (2 out of 12 participants) compared to the running group (0 out of 12 participants).

**Fig. 3:**
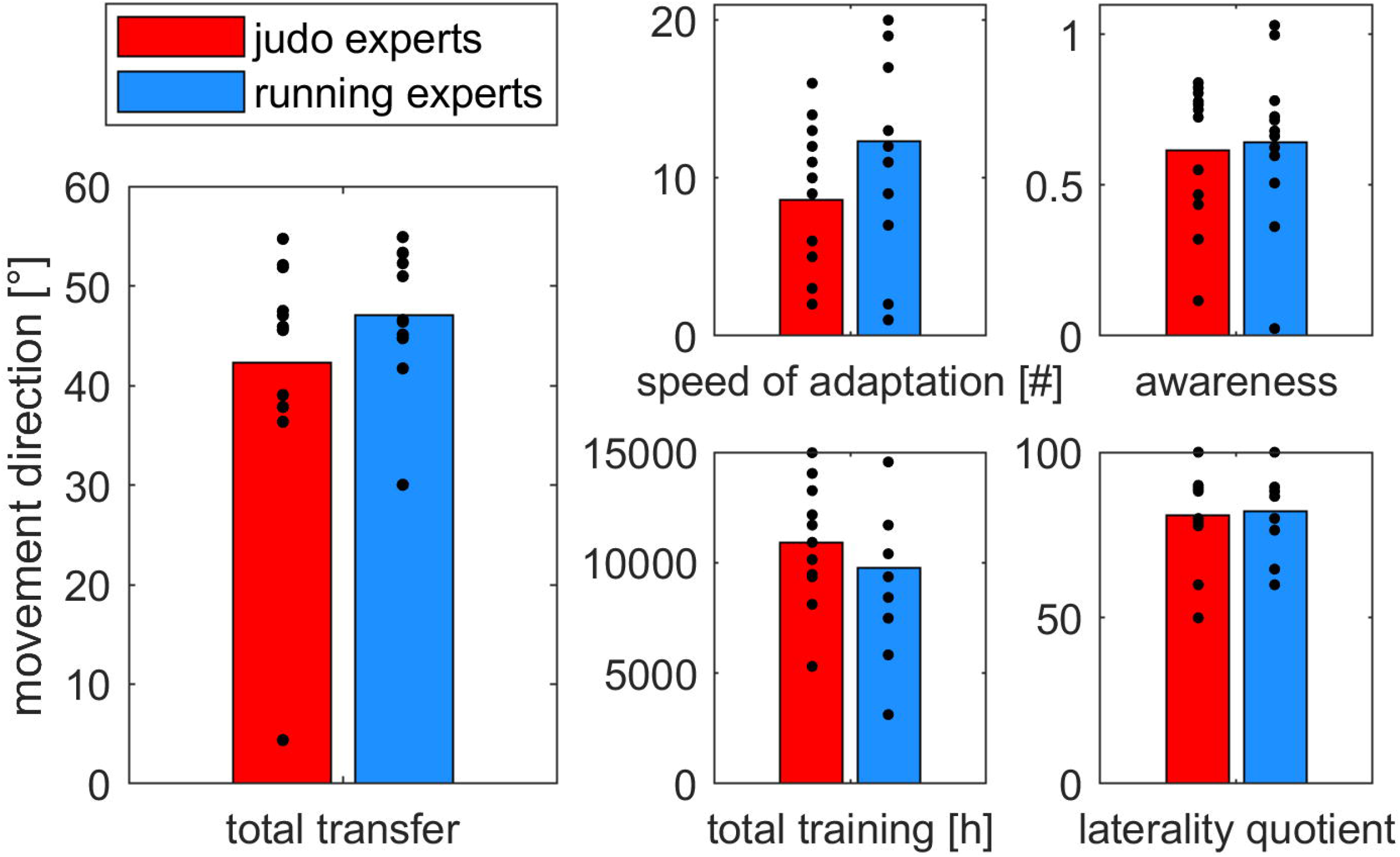
Total intermanual transfer and independent variables. Shown are group mean values for total intermanual transfer, speed of adaptation, awareness, total training and laterality quotient of judo and running experts. Dots represent individual data.

A multiple linear regression was run to predict total intermanual transfer from the independent variables group, speed of adaptation, awareness, total training, and laterality quotient. Our variables significantly predicted total intermanual transfer [F(5, 17) = 4.834, p = 0.006, R^2^ = 0.587] but only awareness added significantly to the prediction [p < 0.001, B = 33.800, beta weight 0.750]. Our estimates therefore predict that for each 0.1 change of index of awareness total intermanual transfer should change by 3.38°. The data included one point of high leverage, however, influence measured as Cook’s distance was not given at this data point. We further found one outliner according to the studentized deleted residuals and repeated the multiple linear regression analysis excluding this participant. Our variables nevertheless predicted the amount of total intermanual transfer [F(5, 16) = 3.744, p = .020, R^2^ = 0.539] and, again, the only variable significantly adding to the prediction was awareness [p = 0.004, B = 19.219, beta weight 0.657]. To find out, whether some handedness items might add to the prediction more than others, we repeated the multiple linear regression analysis using the laterality quotient of personal space. Again, we found a high level of prediction for the whole model [F(5, 17) = 4.655, p = .007, R^2^ = 0.578] but only awareness reached a significant level [p = 0.001, B = 33.416, beta weight 0.741]. In conclusion, our analyses indicate, that the total amount of transfer can be predicted by the amount of awareness but not by any of the other individual factors. Figure 4 depicts the relationship between total intermanual transfer and our different variables in both subject groups.

**Fig. 4:**
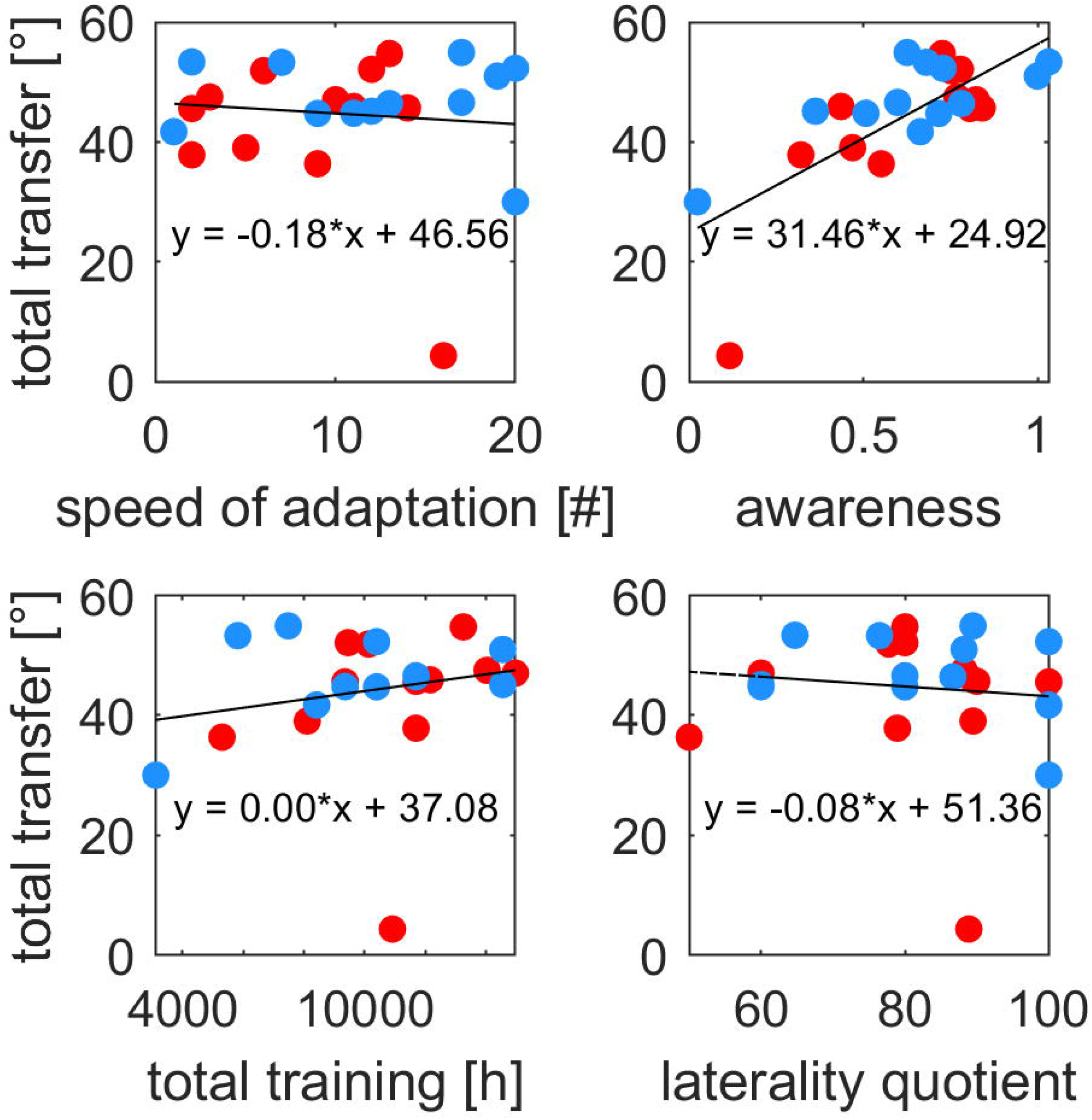
Relationship between total intermanual transfer and independent variables. Shown are the relation between total intermanual transfer and speed of adaptation, awareness, total training or laterality quotient in judo and running experts. Each symbol represents one participant.

Questionnaire data further verified the notion that judo training is mainly bimanual with 96 ± 5 % of the techniques trained including both hands. It further revealed that right-handed judokas mainly use their preferred right side with only 24 ± 21 % of training time is spent with practicing left sided techniques. We aimed at scrutinizing whether the amount of both and left handed training also added to the amount of total intermanual transfer. Multiple linear regression analysis within the judo experts group, however, could not be performed due to significant outliners, influencing points and considerable auto-correlation. Instead we performed Spearman correlations between total intermanual transfer on the one hand and percentage of both or left-handed training on the other hand. Both did not reveal any significant correlations [both: r = −0.433, p = 0.159; left: r = −0.114, p = 0.724].

### 3.3 Implicit intermanual transfer and individual factors

Mean values of implicit intermanual transfer and individual factors of both groups of the second experiment are depicted in Figure 5. Implicit intermanual transfer is very small compared to the total amount of transfer measured in our first experiment (see Fig. 3). Judo athletes show a larger mean transfer (5.2 ± 3.2°) than sports athletes (1.6 ± 4.5°). In accordance, statistical analysis revealed a significant group effect for implicit transfer [t(26) = −2.37, p = 0.026, *d* = −0,930]. Furthermore, we found implicit intermanual transfer of the judo group significantly different from zero [t(11) = −5.56, p < 0.001, *d* = −3,353] but not that of the sports group [t(15) = −1.39, p = 0.184]. There were only four participants in the general sports group reaching implicit transfer values of larger than 5°. Those athletes performed endurance and strength sports as well as racket sports just like the other sports athletes. However, the largest amount of transfer was reached by one gymnast in this group. Across both groups we found a substantial heterogeneity of implicit intermanual transfer values ranging from −7° to 12°. Hence the analysis of inter-individual variations can be a key to uncover the critical predictors driving the amount of implicit transfer (Lefumat *et al.*, 2015).

**Fig. 5:**
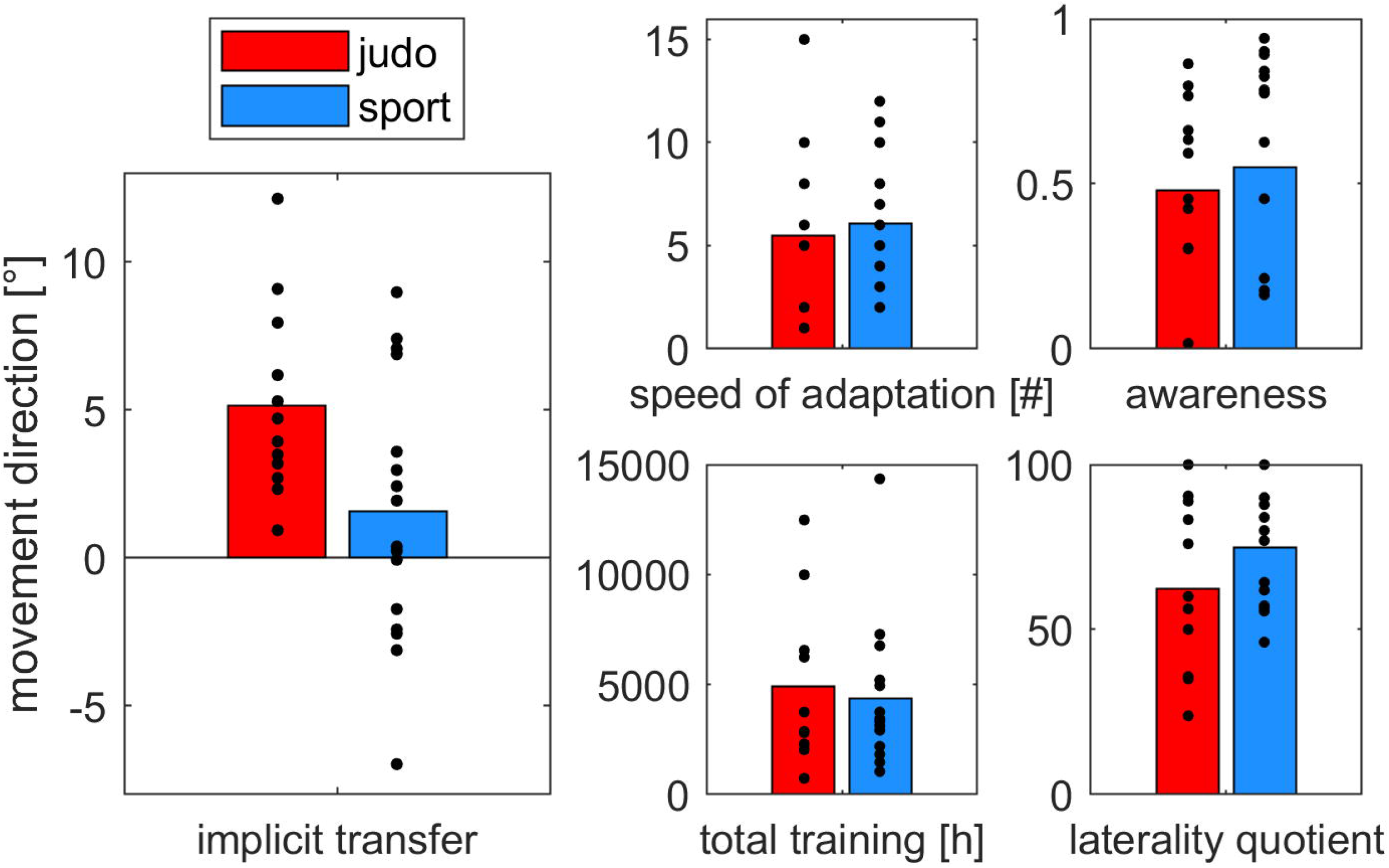
Implicit intermanual transfer and independent variables. Shown are group mean values for implicit intermanual transfer, speed of adaptation, awareness, total training and laterality quotient of judo and sports athletes. Dots represent individual data.

Both groups further show similar values for all individual factors (Fig. 5). Here too, questionnaire data is provided as supporting information S2. The participants of the second experiment were less experienced with total training of around 4600 h compared to those of the first experiment with total training of around 10400 h. Group comparisons within the second experiment revealed no significant differences for speed of adaptation [judo 5.5 ± 4.5; sport 6.1 ± 3.0; t(26) = −0.14, p = 0.888], awareness [judo 0.48 ± 0.30; sport 0.55 ± 0.50; U = −1.07, p = 0.302], total training [judo 4901 ± 3518h; sport 4355 ± 3210h; U = −0.07, p = 0.945] and lateralization quotient [judo 62 ± 25; sport 75 ± 18; t(26) = −0.14, p = 0.888]. Yet as in the first experiment, Edinburgh inventory data show increased both-handedness for items within personal space in the judo group (3 out of 12 participants) compared to the running group (0 out of 16 participants). Together with the (same) result of the first experiment this verifies previous findings of a distinct handedness profile in experienced judokas (Mikheev et al. 2002). Due to our equally trained compare groups we can show for the first time that this change of handedness is specifically related to martial arts and not to general sports training.

To find out which of our parameters contribute to implicit intermanual transfer we again calculated a multiple linear regression with the independent variables group, speed of adaptation, awareness, total training, and laterality quotient. Our variables significantly predicted implicit intermanual transfer [F(5, 22) = 7.694, p < 0.001, R^2^ = 0.636]. All variables but awareness added significantly to the prediction: group [p = 0.003, B = −3.865, beta weight 0.452], speed of adaptation [p = 0.016, B = −0.410, beta weight −0.348], total training [p = 0.002, B= 0.001, beta weight 0.475] and lateralization quotient [p = 0.009, B = 0.078, beta weight 0.392]. Our estimates predict, for example, that for each 1000 hours of total training implicit intermanual transfer should change by 1°. Our estimates also predict that lager lateralization quotient, i.e. stronger right handedness, as well as increased time spent at plateau is related to larger implicit intermanual transfer. Repeating the analysis with the lateralization quotient within personal space yielded the same result [p = 0.034, B = 0.040, beta weight 0.350], also relating stronger right handedness within those items to larger implicit transfer. Standardized coefficients beta weight further show that the amount of total training has the largest effect on implicit intermanual transfer closely followed by the variable group. Our data included several leverage points but none of them were deemed influential by the measure of Cook’s distance. Figure 6 shows the relationship between implicit intermanual transfer and our different variables for judo and sports athletes.

**Fig. 6:**
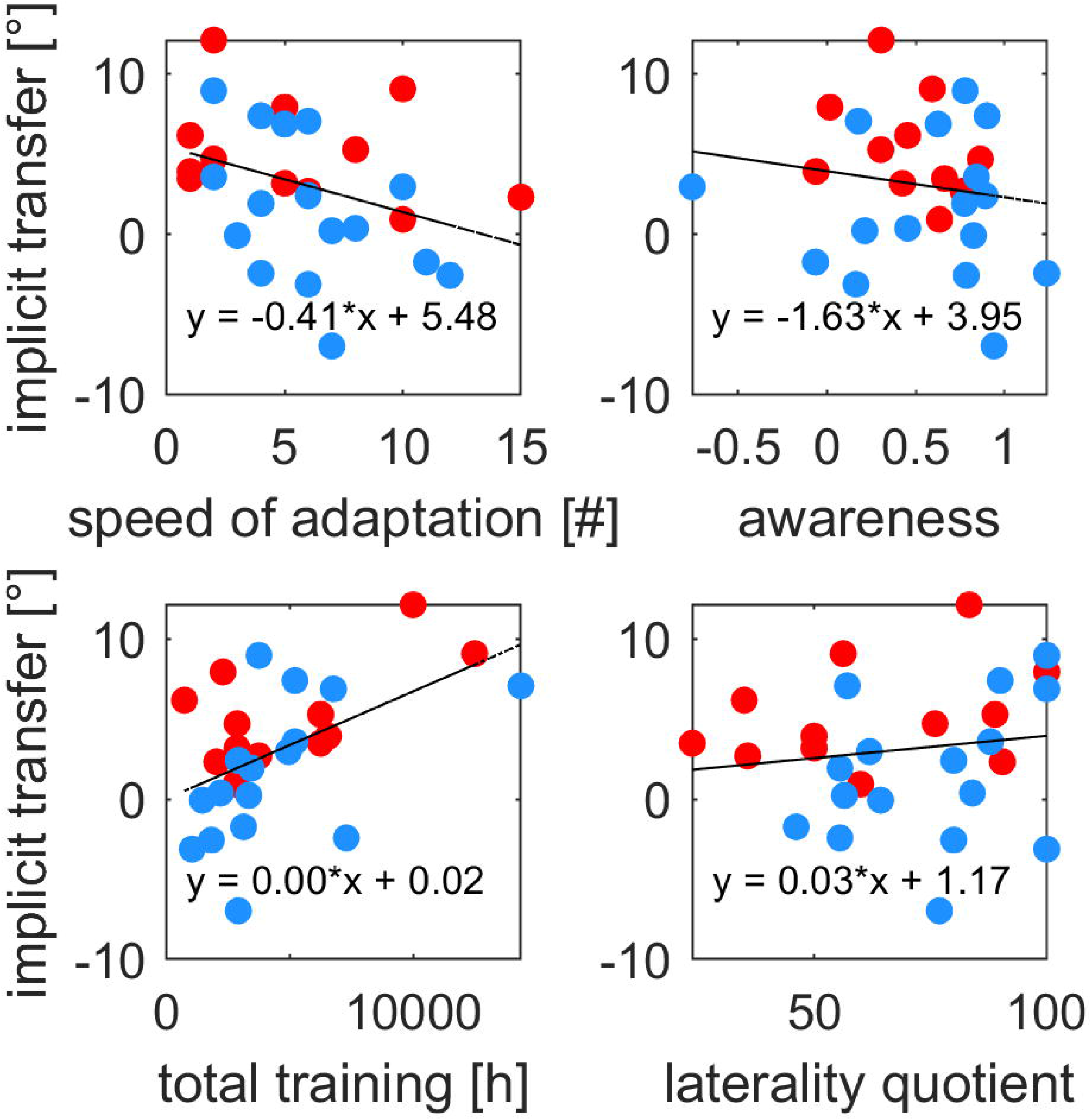
Relationship between implicit intermanual transfer and independent variables. Shown are the relation between implicit intermanual transfer and speed of adaptation, awareness, total training or laterality quotient in judo and sports athletes. Each symbol represents one participant.

Questionnaire data further revealed that 95 ± 5% of practiced techniques include both hands but only 36 ± 14% of training include practice of left sided techniques. We repeated the multiple linear regression analysis within the Judo group adding those two independent variables. Even though the analysis revealed a high correlation with R^2^ = 0.702, number of participants was too small for reaching a significant prediction of implicit intermanual transfer [F(6, 5) = 1.959, p = 0.239]. Only total training revealed a marginally significant prediction [p = 0.059, B = 3.113, beta weight 0.684]. This result suggests that within the judo group total amount of training seems to drive the amount of implicit intermanual transfer more than specific content or other factors such as speed of adaptation, awareness or laterality quotient. This data also included several leverage points but none of them were influential according to Cook’s distance. Repeating the multiple linear regression within the general sports group yielded a significant result [F(4, 11) = 3.652, p = 0.040, R^2^ = 0.570] and only total training significantly added to the prediction within this group [p = 0.049, B = 0.001, beta weight 0.475].

In our experiments we could not directly measure the size of explicit transfer. However, since the total amount of transfer should be the sum of explicit and implicit transfer, we used the data of the second experiment to estimate the amount of explicit intermanual transfer of the first experiment. Therefore, we subtracted the mean group values of implicit intermanual transfer from experiment two from individual transfer data of experiment one and repeated our multiple linear regression analysis. Our variables significantly predicted estimated explicit intermanual transfer [F(5, 17) = 5.637, p = 0.003, R^2^ = 0.624] with group [p = 0.038, B = 8.289, beta weight 0.379] and awareness [p < 0.001, B = 33.800, beta weight 0.716] significantly adding to the prediction. Exclusion of one outliner according to the studentized deleted residuals led to a high auto-correlation of the model according to Durbin-Watson statistic. Note that our calculation of explicit intermanual transfer can only be a rough estimate due to group differences such as sports practiced (running versus general sports), amount of total training (9776 ± 3480h versus 4355 ± 3210h) or age (19 ± 1 versus 26 ± 5). This finding shows that the estimated explicit intermanual transfer can also be predicted by awareness but not by speed of adaptation, laterality quotient or amount of training. The significant effect of group with larger explicit transfer in runners is due to the larger implicit transfer in the judo group of experiment two for one thing. Then again it also relates well to the slightly larger mean awareness index in the running group [running experts 0.64 ± 0.27, judo experts 0.62 ± 0.23], which also implies a larger explicit adaptation process within this group.

## 4. Discussion

The current study examined the impact of individual factors on the amount of intermanual transfer of visuomotor adaptation in judo and sports athletes. We measured the total amount of intermanual transfer (explicit plus implicit) by telling the participants to repeat what was learned during adaptation and the amount of implicit intermanual transfer by instructing the participants to refrain from using what was learned but to perform movements as during baseline. Our main findings show that the total amount of intermanual transfer is related to awareness of the perturbation while implicit intermanual transfer is related to amount of total training, judo training, speed of adaptation and handedness. In addition, we found implicit transfer to be very small compared to total transfer and we found considerable implicit intermanual transfer in judo but not in sports athletes.

### 4.1 Explicit intermanual transfer

Since our implicit intermanual transfer was very small, our total intermanual transfer largely represents the explicit part of transfer. It is thus not surprising, that the only driving factor for total transfer is the amount of awareness of the nature of the perturbation. Former findings about the relation between awareness and intermanual transfer generally point toward an overall picture: Modulation of the amount of awareness by adaptation to either a sudden introduction of the perturbation or a gradual introduction in a stepwise fashion revealed different intermanual transfer after gradual and sudden adaptation in some (Malfait & Ostry, 2004; Michel *et al.*, 2007) but not in other studies (Taylor & Ivry, 2011; Wang, Joshi, *et al.*, 2011; Joiner *et al.*, 2013). We solved this apparent contradiction by our recent work showing larger awareness after adaptation to large than to small perturbations (Werner *et al.*, 2015) and showing larger intermanual transfer after sudden than after gradual adaptation when the perturbation was large but not when it was small (Werner *et al.*, 2019). Furthermore, explicit instructions about the nature of the perturbation lead to an increase of the explicit process of adaptation (Benson *et al.*, 2011) and is also related to an increase of awareness (Werner *et al.*, 2015). Together with the present findings this indicates that paradigms reinforcing the explicit process of adaptation at the same time reinforce the explicit intermanual transfer and both are related to awareness.

In the present study we were not able to directly measure the amount of explicit intermanual transfer. Poh et al. (2016) previously determined explicit and implicit transfer with the help of a reporting method. However, since reporting continued during transfer, the paradigm may have encouraged an explicit transfer as participants made an effort to maintain consistency with the earlier reported strategy. Nevertheless we are confident that explicit intermanual transfer is also related to awareness. There are two reasons: First, implicit intermanual transfer was very small, not even significantly different from zero in our general sports group. Our results for the total amount of transfer (explicit plus implicit), thus, give a good approximation for the amount of explicit intermanual transfer. Second, our estimate of explicit transfer using the data of both experiments also revealed awareness as the only individual factor predicting the amount of transfer. Our findings, therefore, provide converging evidence of the exclusive involvement of awareness in explicit intermanual transfer.

### 4.2 Implicit intermanual transfer

Our multiple linear regression analysis revealed four relevant individual factors driving implicit intermanual transfer: total training, judo training, speed of adaptation and lateralization quotient. The strongest predictor across groups and also within each group was the *total amount of training*. Since we included a wide variety of athletes practicing for example ball sports, racket sports, individual sports or martial arts training, our data suggest that any sports training can lead to increased generalization of motor learning from the dominant to the non-dominant limb. However, noticeable effects can only be expected after around 5,000 h of training which corresponds, for example, to 6 h per week for around 16 years (see Fig. 6). Apart from general sports practice, *judo training* further proved to be a crucial factor driving the amount of implicit intermanual transfer. We are aware that the direction of causality should be specified here. Does judo training, in particular, lead to improved implicit intermanual transfer or does this respective ability predispose a person to choose judo as a sport? At first glance, our data supports the second possibility since all sports training lead to increased implicit transfer and judokas generally show a higher level of transfer. However, it is also possible that initial judo training – our participants all had at least seven years of training (see supporting information S2) – is particularly beneficial for the ability to generalize motor learning. Moreover, judo specific hemispheric lateralization is a result of long-time judo training as argued by Mikheev et al. (Mikheev *et al.*, 2002): Results of different lateralization tasks were fine-grained in judokas and interrelated specifically with judo characteristics. For example, a number of handedness factors dealing with objects in peripersonal space – such as a judo opponent in near space – as well as wrestling stand side correspond with respect to lateralization. Furthermore, in their study the stronger, pushing leg was never used for wagging in their control participants, but there was a relationship between stronger and wagging leg in their judokas. This also suggests a specific training effect since the stronger leg can be a good choice for a wag in judo techniques. A third individual factor predicting the amount of implicit intermanual transfer is the *lateralization quotient*. The stronger right-handed our participants were the larger was their implicit transfer. This is consistent with the results of a previous study showing a positive correlation between right or left-handedness and intermanual transfer values (Lefumat *et al.*, 2015). In contrast, Chase and Seidler (2008) found an increase of intermanual transfer in relation to a decrease of the degree of handedness. However, this was only true for left-handed but not for right-handed individuals. In addition, both directions of transfer (dominant hand to non-dominant hand and vice versa) were measured in that study and combined results were correlated to lateralization quotients. The last significant individual factor uncovered by our findings was the *speed of adaptation*. This finding is in line with previous studies on dynamic motor adaptation reporting a more stable intermanual transfer after long than after short sudden training (Joiner *et al.*, 2013) and greater intermanual transfer after extended time spent at plateau (Block & Celnik, 2013). Taken together, our findings show for the first time that sports training enhances generalization of motor learning and that judo training, in particular, is related to larger intermanual transfer. Moreover, on the basis of the present results we can specify that previously reported individual factors such as speed of adaptation and laterality quotient are particularly related to implicit intermanual transfer.

### 4.3 Neural correlates and mechanisms of intermanual transfer

Studying the way in which motor learning generalizes is a very powerful method for understanding the underlying neural processes (Shadmehr, 2004). Wang et al. (2015) suggested that intermanual transfer of sensorimotor adaptation might involve several aspects of a neural representation, some of which are effector dependent and others are effector independent. There is also neuropsychological evidence (from a serial button-press motor task) indicating different neural networks for explicit and implicit transfer of learning (Jung *et al.*, 2019). It could be speculated that the implicit intermanual transfer is effector dependent and the explicit intermanual transfer is effector independent. We suggest that the explicit process of adaptation measured by awareness does not directly relate to a motor action of a specific limb and that it might, thus, relate to an effector independent neural representation. Since the explicit process plays a role in early adaptation (Benson *et al.*, 2011) and early learning was found to engage prefrontal brain regions (e.g. Clower et al. 1996; Anguera et al. 2007; Inoue et al. 2015), especially the dorsolateral prefrontal cortex (Anguera *et al.*, 2011), this brain area could be associated with the explicit process of adaptation (Taylor & Ivry, 2014) and, consequently, with explicit intermanual transfer. This argument is supported by the fact that the explicit process of adaptation can be entirely transferred to the other hand (Poh *et al.*, 2016).

The implicit process, then again, is only partly accessible by the other limb (Poh *et al.*, 2016) supporting the notion of an effector dependent neural representation. Nevertheless, the distinct neuronal mechanisms underlying (implicit) intermanual transfer have long been discussed and are still an open question today (Joiner *et al.*, 2013). Two general positions can be identified: Transfer either results from two separate internal models in each hemisphere updated through callossal connections or results from one internal model located in the contralateral hemisphere and ipsilateral projections to the untrained hand (Criscimagna-Hemminger *et al.*, 2003). Our findings have the potential to shed new light on this ongoing debate.

On the one hand we found enhanced implicit intermanual transfer in judo athletes. Decreased hemispheric lateralization in judokas (Mikheev *et al.*, 2002) might have resulted in greater involvement of the ipsilateral hemisphere during learning. This is in line with the idea of two separate internal models for each limb in both hemispheres that are updated during adaptation (Taylor & Heilman, 1980; Parlow & Kinsbourne, 1989; Thut *et al.*, 1996; Sainburg & Wang, 2002). The mechanism of this process seems to be a decrease of transcallosal inhibition (Cook, 1986) during training which leads to larger right hemispheric motor control than under untrained conditions (Mikheev *et al.*, 2002; Perez *et al.*, 2007; Chase & Seidler, 2008) and the subsequent storage of over-learned internal models in both hemispheres. It can be speculated that our judo athletes show an even larger decrease of inter-hemispheric inhibition (Mikheev *et al.*, 2002) than our sports athletes. In fact, well-trained musicians showed decreased transcallosal inhibition in a transcranial magnetic stimulation study (Ridding *et al.*, 2000) compared to control participants. The notion of dual internal models is substantiated by findings of bilateral activation of several brain regions during adaptation such as primary and secondary somatosensory cortex (Diedrichsen *et al.*, 2005), deep nuclei of the cerebellum and prefrontal cortex (Nezafat *et al.*, 2001). Further support for this model comes from research on intermanual transfer in split-brain and acallosal patients. Several studies show impaired intermanual transfer of a variety of unimanually learned motor skills (Ferriss & Dorsen, 1975; McQ. Reynolds & Jeeves, 1977; Gott & Saul, 1978; Jeeves, 1979; De Guise *et al.*, 1999) including visuomotor adaptation (Lassonde *et al.*, 1995; Bao *et al.*, 2018) and imply that transfer depends on intact interhemispheric communication via the corpus callosum. Contrastingly, one recent study did show intermanual transfer of adaptation to a force field in a split-brain patient (Criscimagna-Hemminger *et al.*, 2003), however, learning instead of transfer was presumably measured there (Joiner *et al.*, 2013).

On the other hand we found enhanced implicit intermanual transfer in stronger right handed participants. Stronger right-handedness might have resulted in greater activation of the contralateral left hemisphere during adaptation with the right arm (Dassonville *et al.*, 1997; Pool *et al.*, 2014) and greater involvement of the ipsilateral left hemisphere when using the left hand during intermanual transfer (Verstynen *et al.*, 2005). This is in line with the idea that the neural representation within the trained contralateral hemisphere results in intermanual transfer through ipsilateral projections. Support for this model comes from the fact that damage to the dominant left hemisphere can impair the motor function of both right and left hands (Haaland & Harrington, 1996; Sainburg, 2014) and sparing of left hand motor control following callosal or left hemispheric lesions (Geschwind & Kaplan, 1962; Kertesz *et al.*, 1984). In addition, there are a small but meaningful number of corticospinal projections to proximal arm muscles from the ipsilateral hemisphere (Galea & Darian-Smith, 1997) and the dominant hemisphere might play an important role in controlling the non-dominant limb via ipsilateral projections (Haaland & Harrington, 1996). Within this framework intermanual transfer does not rely on interhemispheric transfer of information across the body of the corpus callosum.

To sum up, increased implicit intermanual transfer in judo athletes is in line with the idea of two separate internal models in each hemisphere while increased implicit transfer in strongly right handed participants is in line with the idea of a strong neural representation in the contralateral hemisphere and ipsilateral projections during transfer. Moreover, there has been converging neurophysiological evidence for both models. Instead of distinguishing between those options our data underpins the possibility that both are possible neural mechanisms and that they could be applied according to individual predisposition.

Given that we only tested intermanual transfer from the right, dominant to the left, non-dominant arm our results are restricted to this direction of transfer. While asymmetry of intermanual transfer of motor adaptation was first proposed by Sainburg and Wang (Sainburg & Wang, 2002) an heterogeneous picture has emerged since then: Some studies found transfer only from the dominant to the non-dominant arm after dynamic adaptation (Criscimagna-Hemminger *et al.*, 2003; Wang & Sainburg, 2004) but others found equal transfer in both directions (Wang, Przybyla, *et al.*, 2011; Stockinger *et al.*, 2015). Importantly, Poh et al. (2016) neither found an asymmetry of total transfer nor of explicit transfer. This further supports our notion of an effector independent neuronal representation of explicit transfer. Implicit intermanual transfer, however, could be differentially related to handedness. While our results are in line with previous findings of larger transfer in strong right-handedness (Lefumat *et al.*, 2015) there are conflicting results for strong left-handedness (Chase & Seidler, 2008; Lefumat *et al.*, 2015) indicating different neuronal mechanisms for strongly right or left handed individuals (Cherbuin & Brinkman, 2006a, 2006b).

### 4.4 Amount of intermanual transfer

In the present study we found a total amount of intermanual transfer of 72 % in judo and of 81 % in running experts compared to final adaptation level (as mean of last three adaptation episodes). Former work shows a wide span of intermanual transfer values from the dominant right to the non-dominant left arm ranging from no transfer at all (Martin *et al.*, 1996; Kitazawa *et al.*, 1997) up to 75 % of final adaptation level (Poh *et al.*, 2016). Generally, the amount of intermanual transfer of sensorimotor adaptation is rather limited (Joiner *et al.*, 2013) reaching around 10-45 % (e.g. Wang and Sainburg 2003, 2006; Chase and Seidler 2008; Taylor et al. 2011; Joiner et al. 2013; Lefumat et al. 2015). No specific instructions during transfer tests were given in any of these studies, suggesting that all of them measured the total amount of intermanual transfer. If the explicit process of adaptation is reduced due to small perturbation size (Werner *et al.*, 2015) or gradual adaptation paradigms (e.g. Wang et al. 2011) this probably also leads to a reduced amount of explicit intermanual transfer. Given the low proportion of implicit transfer in our data this should result in a rather limited total amount of intermanual transfer. Therefore, our findings indicate that different proportions of implicit and explicit intermanual transfer were previously determined and this could be one explanation for the large range of reported intermanual transfer values.

So far only one previous study differentiated between explicit and implicit transfer and found an implicit intermanual transfer of 25 % compared to final adaptation level (Poh *et al.*, 2016). Our results conversely show an implicit transfer of 14 % in judo and of 2 % in sport athletes compared to final adaptation level. This discrepancy could be explained by the fact that Poh et al. measured transfer over several time points during adaptation phase which might have induced actual learning with the (untrained) limb causing increased implicit transfer (Huberdeau *et al.*, 2015). Our findings emphasize the importance of instructions before intermanual transfer tests because they determine whether the implicit or the total amount of transfer is measured. Furthermore, transfer should best be assessed at the end of adaptation to prevent confounding transfer and learning effects.

### 4.5 Generalization of results

It can be argued that our results only hold for athletes. Even our control participants were either highly experienced runners competing on a national or even international level in the first study or sports athletes with at least four years of training in their corresponding sports in the second study. However, we assume that our findings can be generalized to a broader population since all individual factors have previously been reported to drive intermanual transfer: Handedness scores (Chase & Seidler, 2008; Lefumat *et al.*, 2015), time spent at plateau (Block & Celnik, 2013; Joiner *et al.*, 2013) and awareness (Malfait & Ostry, 2004; Poh *et al.*, 2016; Werner *et al.*, 2019) all predicted the amount of intermanual transfer in non-specialized populations.

In summary, the present findings demonstrate that different individual factors differentially relate to explicit and implicit intermanual transfer. Specifically, explicit transfer can be strongly predicted by awareness of the learned perturbation while implicit transfer is related to judo training but also to the total amount of training, time spend at plateau and handedness. Our results indicate that paradigms reinforcing the explicit process of adaptation at the same time reinforce explicit intermanual transfer and both are related to awareness. Furthermore, our findings show for the first time that sports training enhance generalization of motor learning and that judo training, in particular, is related to larger intermanual transfer. On the basis of the present results we can further specify that previously reported individual factors such as speed of adaptation and handedness are particularly related to implicit but not to explicit intermanual transfer. Finally we speculate that implicit intermanual transfer involves effector dependent while explicit intermanual transfer involves effector independent neural representations.

## Acknowledgments

This work was supported by a DAAD Travel Grant 91721115 awarded to Susen Werner as well as a DAAD Partnership Program Grant 57320531 awarded to Tobias Vogt. Furthermore, we would like to thank Prof. Opher Donchin for helpful comments and discussions about our data.

## Author Contributions

SW, KK and TV contributed to the idea and design of the experiment. SW, KK and TG carried out the data collection and processed the experimental data. SW drafted the manuscript and designed the figures. All authors discussed the results and contributed to the final manuscript.

## Data Accessibility Statement

All data are available in the online repository (https://osf.io/khgzs/).

